# Median Preoptic *Brs3*-Expressing Neurons Regulate Heat-Evoked Defensive Escalation

**DOI:** 10.64898/2026.09.28.755097

**Authors:** Zixu Zhang, Shengru Hu, Hongchao Cao, Wei Xie, Zhiping Cao, Mingdao Mu

## Abstract

Animals confronted with increasing heat must switch from exploratory behavior to vigorous actions that limit further exposure, but the forebrain circuits controlling this transition are poorly understood. We exposed male mice to progressively stronger floor heating while providing continuous access to a climbing mesh separated from the heating plate. Higher plate settings shifted behavior from rearing toward wall-directed digging and climbing, with jumping at the highest setting, and increased c-Fos labeling in the median preoptic nucleus. Neurons expressing bombesin receptor subtype 3 showed greater c-Fos recruitment under strong heating, and their population calcium signals increased when mice entered the plate-controlled zone and decreased when they left. Optogenetic activation without heating produced place avoidance, vigorous actions, and increased locomotion. Inhibition reduced locomotion and, during strong heating, shifted behavior from climbing and jumping toward rearing and weakened avoidance of a localized heat source. These findings identify median preoptic *Brs3*-expressing neurons as regulators of behavioral escalation and heat avoidance.

## Introduction

Heat elicits a range of protective responses, from rapid withdrawal to spatial avoidance and escape (Barik et al., 2018; Harikai et al., 2004; Jung et al., 2022). Which response is expressed depends on both stimulus intensity and the opportunities available to reduce further exposure (Almeida et al., 2015; Harikai et al., 2004; Jung et al., 2022). Peripheral heat-sensitive receptors and the spinal and brainstem pathways that generate rapid nocifensive responses are comparatively well defined (Barik et al., 2018; Vriens et al., 2011). How forebrain circuits reorganize behavior toward more vigorous actions as heating intensifies remains less clear (Carta et al., 2026; Grajales-Reyes et al., 2024; Jung et al., 2022).

The preoptic area (POA) receives cutaneous thermosensory information relayed through the lateral parabrachial nucleus (LPB) and coordinates physiological and behavioral responses to thermal challenge (Morrison and Nakamura, 2019; Nakamura, 2018; Yang et al., 2020; Zhao et al., 2017). Molecularly defined POA populations regulate heat loss, thermogenesis, cardiovascular output, and behavioral temperature selection, highlighting the functional diversity of this region (Machado and Saper, 2022; Norris et al., 2021; Piñol et al., 2021). Within the POA, the median preoptic nucleus (MnPO) is a principal recipient of ascending thermosensory projections from the LPB (McKinley et al., 2021; McKinley et al., 2015; Nakamura, 2018; Yahiro et al., 2023). Activation of glutamatergic MnPO neurons promotes cutaneous vasodilation, whereas LPB-to-MnPO transmission mediates avoidance of a heated surface (Abbott and Saper, 2017; Yahiro et al., 2023). These findings establish the MnPO as a site at which ascending thermal information can engage both physiological heat loss and behavioral heat avoidance (Zhou et al., 2023). Whether molecularly defined MnPO populations control the transition toward vigorous defensive actions during stronger heating remains unresolved.

The MnPO is molecularly heterogeneous and participates in body-fluid homeostasis as well as thermal regulation (Machado and Saper, 2022; McKinley et al., 2015; Yao et al., 2023). One MnPO population expresses *Brs3*, which encodes bombesin receptor subtype 3, an orphan G protein-coupled receptor (Mogul et al., 2021; Xiao and Reitman, 2016; Yao et al., 2023). *Brs3*-expressing neurons in the hypothalamus and POA regulate body temperature, energy expenditure, and cardiovascular function (Lateef et al., 2014; Piñol et al., 2021; Piñol et al., 2018; Xiao et al., 2017). Previous studies targeting a broad POA *Brs3*-expressing population that encompassed the MnPO and adjacent preoptic regions showed that its manipulation altered body temperature, heart rate, and blood pressure (Piñol et al., 2021). This work established a role for POA *Brs3*-expressing neurons in physiological thermal regulation, particularly cold defense, but did not isolate the MnPO-centered population or examine its contribution to active defensive behavior during strong heating.

Here, we asked whether MnPO *Brs3*-expressing neurons shape the transition toward vigorous defensive actions as floor heating intensifies. Using a floor heating paradigm in which mice could climb away from the heated surface, we combined c-Fos mapping and fiber photometry with bidirectional optogenetics and complementary behavioral assays. We found that MnPO *Brs3*-expressing neurons were increasingly recruited under stronger floor heating and contributed to place avoidance, vigorous behavioral output, and avoidance of a localized heat source. These findings identify a molecularly defined MnPO population that links thermal challenge to defensive escalation.

## Results and Discussion

### Stronger Floor Heating Shifts Behavior Toward Vigorous Defensive Actions and Increases MnPO c-Fos Labeling

We first examined how the behavioral repertoire changed during 5 min floor heating sessions. Separate cohorts were tested with the heating plate set to 25, 35, 45, or 55°C, while a wall-mounted mesh remained accessible throughout each session (Fig 1A). The behavioral repertoire changed as the plate setting increased (Fig 1B). Rearing predominated at 25 and 35°C, whereas wall-digging and climbing became more prominent at the higher settings (Fig 1C–N). Jumping emerged at 55°C, where both its total duration and event number were greater than at 25°C (P < 0.001; Fig 1O–R). Wall-digging was directed toward the chamber boundary, climbing transferred mice from the acrylic floor to the wall-mounted mesh, and jumping briefly removed all four paws from the floor. Increasing the plate setting therefore shifted behavior from rearing toward more vigorous actions, with climbing and jumping directly reducing contact with the heated surface.

**Figure 1.**
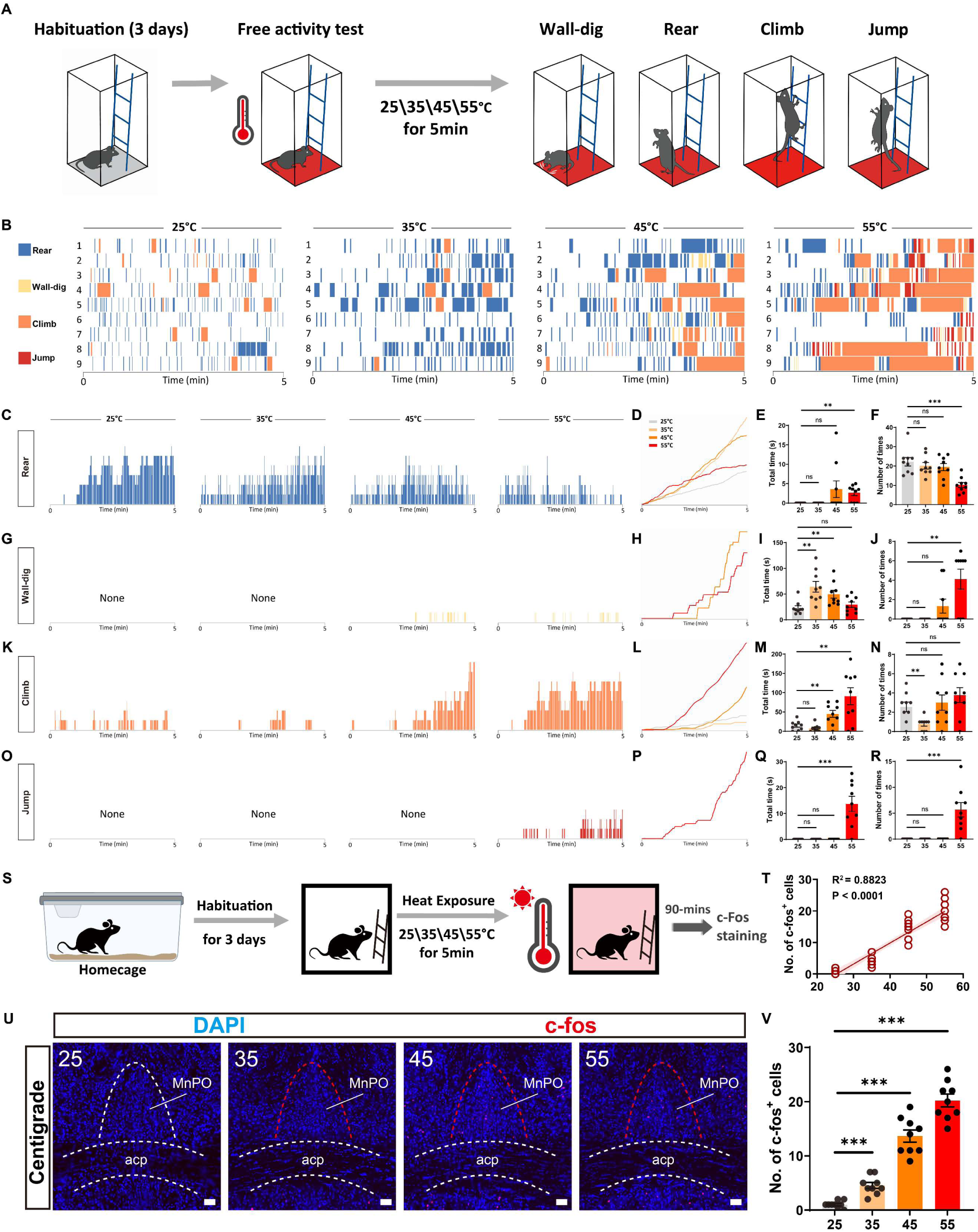
Increasing floor heating shifts behavior toward vigorous defensive actions and increases MnPO c-Fos labeling. (A) Experimental design of the floor heating assay. After 3 days of habituation, separate cohorts of mice were tested for 5 min in a chamber with an acrylic floor 5 mm thick positioned above a temperature-controlled heating plate set to 25, 35, 45, or 55°C. The indicated temperatures denote heating-plate settings and do not represent temperatures measured at the acrylic surface or paw interface. A wall-mounted mesh insert, physically separated from the heating plate, remained accessible throughout the session and permitted mice to leave the acrylic floor by climbing. (B) Ethograms from individual mice tested at each plate setting. Each row represents one mouse (n = 9 mice per condition). (C–F) Rearing, displayed as event rasters, cumulative duration, total duration, and event count. (G–J) Wall-digging, displayed in the same sequence. (K–N) Climbing. (O–R) Jumping. For (C–R), n = 9 biologically independent mice per condition. (S) Experimental design for regional c-Fos mapping. After 3 days of habituation, mice were exposed to one heating-plate setting for 5 min and perfused 90 min after the beginning of the session. (T) Relationship between heating-plate setting and the number of c-Fos-positive cells in the sampled MnPO region (R² = 0.8823, P < 0.0001, linear regression). Each point represents one section. Nine anatomically matched sections from three mice were analyzed per condition. Sections obtained from the same mouse represent nested observations. (U) Representative c-Fos immunostaining with 4’,6-diamidino-2-phenylindole (DAPI) counterstaining at each plate setting. acp, posterior part of the anterior commissure. Scale bar, 100 µm. (V) Quantification of MnPO c-Fos-positive cells across heating-plate settings. Behavioral analyses used one value per mouse. Behavioral measures and section-level c-Fos counts at 35, 45, and 55°C were each compared with the corresponding 25°C condition using two-tailed unpaired Student’s t-tests. Bar graphs show mean ± SEM. *P < 0.05; **P < 0.01; ***P < 0.001; ns, not significant.

Strong heating is known to recruit jumping and other escape-like responses through nocifensive circuits operating at peripheral, spinal, brainstem, and forebrain levels (Barik et al., 2018; Harikai et al., 2004; Jung et al., 2022). Here, defensive escalation describes the redistribution of behavior toward more vigorous actions within this assay. It does not imply that rearing, wall-digging, climbing, and jumping form a fixed hierarchy that applies across defensive contexts.

We next assessed MnPO c-Fos expression after exposure to the same floor heating conditions (Fig 1S–V). Across nine sections obtained from three mice per condition, MnPO c-Fos-positive cell counts increased with the heating-plate setting (section-level regression: R² = 0.8823, P < 0.0001; Fig 1T,V). The transition toward more vigorous heat-evoked behavior was therefore accompanied by increasing c-Fos labeling in the sampled MnPO region.

### MnPO Brs3-Expressing Neurons Are Recruited as Floor Heating Intensifies

Analysis of publicly available multiplexed error-robust fluorescence in situ hybridization data from the Allen Brain Cell Atlas showed *Brs3*-expressing cells across multiple rostrocaudal levels of the MnPO (Fig 2A). We used Cre-dependent viral vectors in *Brs3*-P2A-iCre mice to examine this population (Fig S1).

**Figure 2.**
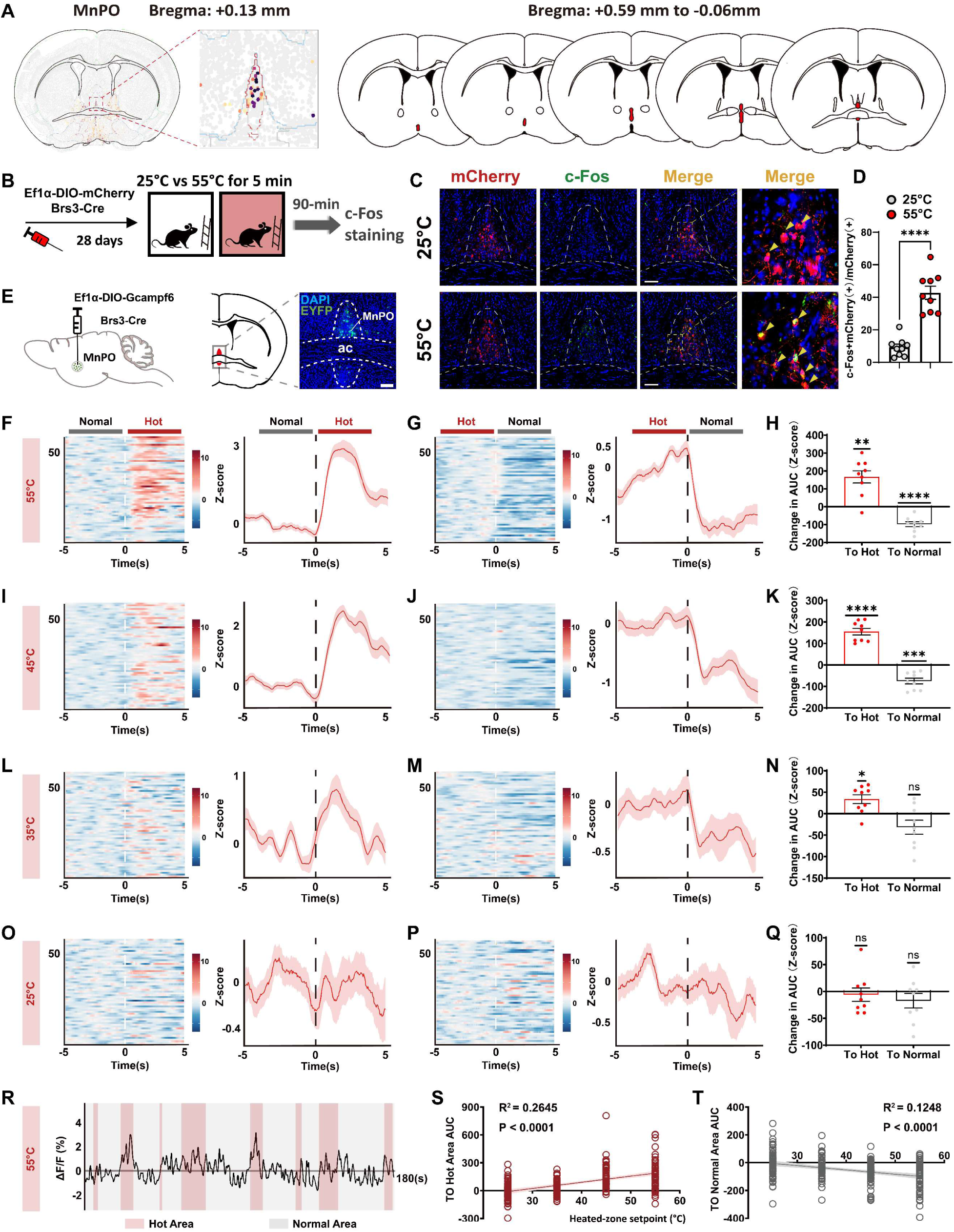
MnPO *Brs3*-expressing neurons are recruited as floor heating intensifies. (A) Spatial distribution of *Brs3*-expressing cells in the median preoptic nucleus (MnPO), visualized using publicly available multiplexed error-robust fluorescence in situ hybridization (MERFISH) data from the Allen Brain Cell Atlas. A representative coronal section at Bregma +0.13 mm and the rostrocaudal distribution of *Brs3*-expressing cells from Bregma +0.59 to −0.06 mm are shown. (B) Experimental design for c-Fos mapping in MnPO *Brs3*-expressing neurons. *Brs3*-P2A-iCre mice received a Cre-dependent adenoassociated viral vector expressing mCherry in the MnPO and were subsequently exposed for 5 min to a heating plate set to either 25 or 55°C. Mice were perfused 90 min after the beginning of the session. (C) Representative images of mCherry and c-Fos labeling after exposure to the two plate settings. Arrowheads indicate mCherry-positive cells containing c-Fos-positive nuclei. Scale bar, 100 µm. (D) Percentage of mCherry-positive cells coexpressing c-Fos. Each point represents one section. Nine anatomically matched sections from three mice were analyzed per condition, with sections from the same mouse representing nested observations. (E) Fiber photometry strategy and representative GCaMP6f expression and optical-fiber placement in the MnPO. ac, anterior commissure. Scale bar, 100 µm. (F, G) Event-aligned heat maps and mean calcium responses during entry into the plate-controlled zone (F) and return to the comparison zone (G) with the plate set to 55°C. (H) Area under the curve (AUC) of the event-associated fluorescence responses during the 5 s following entry and return at the 55°C setting. (I– K) Corresponding entry, return, and AUC analyses at 45°C. (L–N) Corresponding analyses at 35°C. (O– Q) Corresponding analyses at 25°C. ΔF/F was calculated relative to the mean fluorescence during the 2 s preceding each transition. Peri-event traces are displayed as z scores from −5 to +5 s, and AUC was calculated over the 0 to 5 s interval after each transition. At each plate setting, nine mice contributed a total of 58 zone-transition trials, with each mouse contributing six or seven trials. Each trial comprised an entry into the plate-controlled zone and a corresponding return to the comparison zone. Thus, 58 entry events and 58 return events were analyzed at each plate setting. Events obtained from the same mouse represent repeated within-mouse observations. (R) Representative ΔF/F trace during repeated transitions between the comparison and plate-controlled zones at the 55°C setting. (S, T) Relationships between heating-plate setting and event-level AUC during entry into the plate-controlled zone (S; R² = 0.2645, P < 0.0001) and return to the comparison zone (T; R² = 0.1248, P < 0.0001). Linear regression was used for (S, T). A two-tailed unpaired Student’s t test was used for the section-level comparison in (D), and twotailed one-sample t tests against zero were used for the event-level analyses in (H, K, N, Q). Mean traces are shown with SEM shading, and bar graphs show mean ± SEM. *P < 0.05; **P < 0.01; ***P < 0.001; ns, not significant.

To assess neuronal recruitment, we expressed Cre-dependent mCherry in the MnPO and quantified c-Fos after 5 min floor heating with the plate set to either 25 or 55°C (Fig 2B,C). The proportion of mCherry-labeled cells containing c-Fos-positive nuclei was greater at 55°C than at 25°C (nine sections from three mice per condition; section-level comparison, P < 0.0001; Fig 2D). MnPO *Brs3*-expressing neurons were therefore more strongly recruited under the highest heating condition.

For fiber photometry, we expressed Cre-dependent GCaMP6f in the MnPO and implanted an optical fiber above the labeled territory. Post hoc histology localized the viral expression centers to the MnPO and confirmed that the optical-fiber tips were positioned above the labeled cells (Fig 2E). Viral labeling encompassed dorsal and ventral portions of the MnPO, and we therefore refer to the recorded population collectively as MnPO *Brs3*-expressing neurons.

We next monitored population GCaMP6f signals while mice moved voluntarily between a comparison zone and a plate-controlled test zone. With the plate set to 55°C, entry into the test zone was accompanied by a rapid increase in GCaMP6f signal, whereas return to the comparison zone was accompanied by a decrease relative to the 2 s pre-transition baseline (P < 0.0001; Fig 2F–H). Responses changed in the same direction at 45 and 35°C, whereas transitions at 25°C produced no significant change (Fig 2I–Q). A continuous recording at the 55°C setting showed repeated increases and decreases in GCaMP6f signal as the mouse moved between the two zones (Fig 2R).

At each plate setting, nine mice contributed 58 entry events and 58 corresponding return events, with each mouse contributing six or seven repeated trials. In event-level analyses across the four plate settings, entry-associated responses increased with the heating-plate setting (R² = 0.2645, P < 0.0001), whereas return-associated responses became more negative as the setting increased (R² = 0.1248, P < 0.0001; Fig 2S,T). The c-Fos and photometry data thus show greater recruitment of MnPO *Brs3*-expressing neurons under stronger floor heating and rapid changes in their population activity as mice enter and leave the plate-controlled zone.

The MnPO receives ascending thermosensory information from the lateral parabrachial nucleus, and its glutamatergic neurons promote heat loss through cutaneous vasodilation (Abbott and Saper, 2017; Nakamura, 2018; Yahiro et al., 2023). Transmission from the lateral parabrachial nucleus to the MnPO also contributes to avoidance of heated surfaces (Yahiro et al., 2023). Our findings identify *Brs3*-expressing neurons as an MnPO population recruited during strong floor heating and extend the established role of this region from physiological heat loss and spatial heat avoidance to the organization of vigorous defensive behavior.

This behavioral role complements previous studies of *Brs3*-expressing neurons in the broader preoptic area and hypothalamus. Broad manipulation of preoptic *Brs3*-expressing neurons alters body temperature, heart rate, and blood pressure, whereas dorsomedial hypothalamic *Brs3*-expressing neurons regulate thermogenesis, energy expenditure, and cardiovascular output (Piñol et al., 2021; Piñol et al., 2018).

Those studies primarily established physiological functions in cold defense and thermal feedback regulation. The present results reveal a behavioral contribution of an MnPO-centered population during strong heating. Whether the same MnPO *Brs3*-expressing cells coordinate behavioral activation with autonomic thermal responses remains unknown.

The c-Fos and photometry experiments capture different temporal features of neuronal recruitment. c-Fos labeling reflected activity associated with the complete heating session, whereas calcium signals changed rapidly during transitions between zones. These transition-linked signals should not be interpreted as a direct neural representation of temperature alone because zone entry and return also changed movement, posture, floor contact, and the behavioral significance of the animal’s location. Cellular-resolution recordings combined with detailed tracking will be required to determine whether thermal input, aversive value, and preparation for action are represented by shared or distinct MnPO *Brs3*-expressing neurons.

### Activation of MnPO Brs3-Expressing Neurons Produces Place Avoidance and Elicits Vigorous Actions

We expressed Cre-dependent ChR2 in MnPO *Brs3*-expressing neurons (Fig 3A). In a real-time place preference assay, ChR2-expressing mice spent less time in the photostimulation-paired chamber than controls (P < 0.01; Fig 3B,C). Activation of this population was therefore sufficient to produce place avoidance.

**Figure 3.**
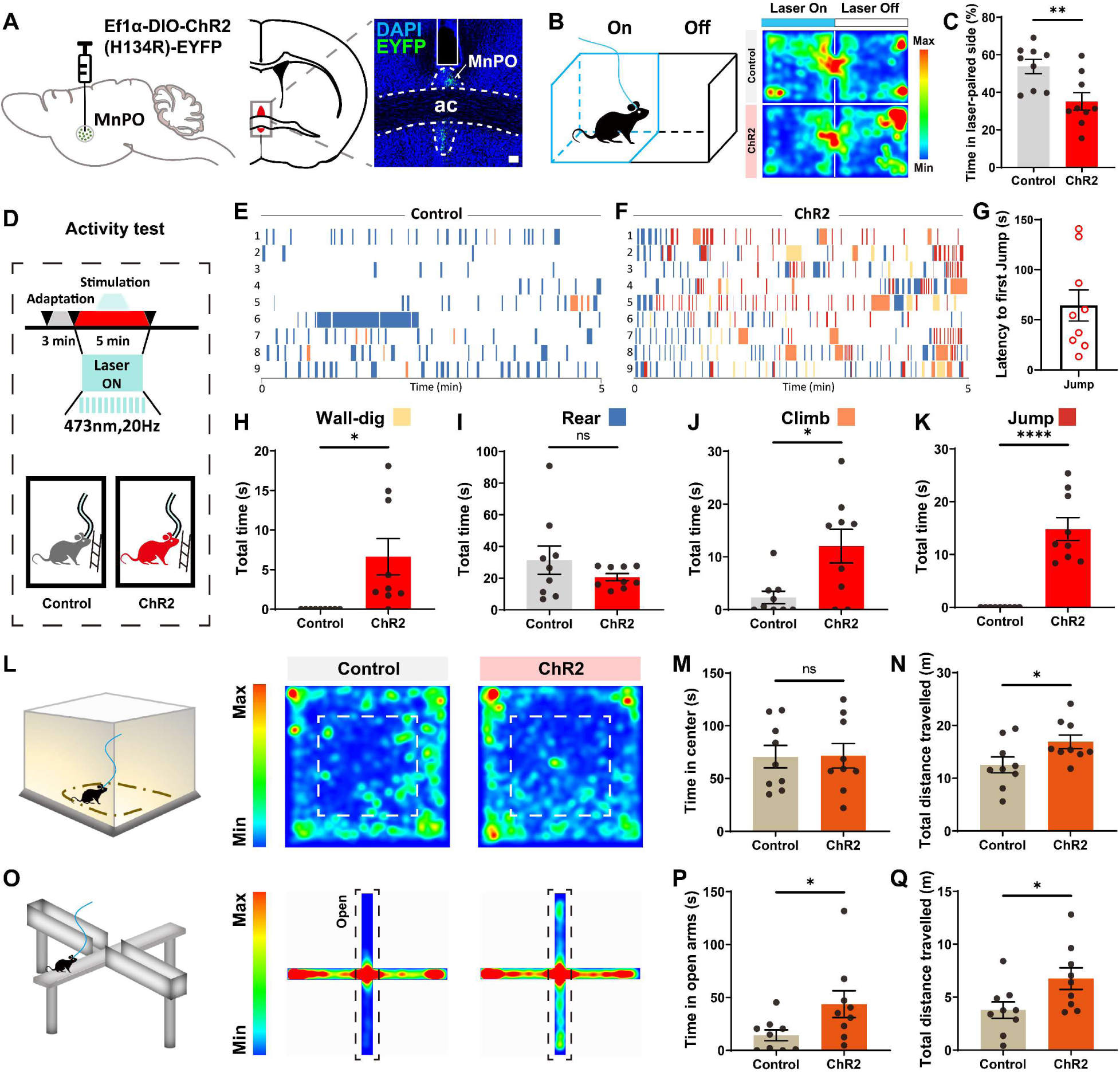
Activation of MnPO *Brs3*-expressing neurons produces place avoidance and elicits vigorous actions. (A) Experimental strategy for optogenetic activation of MnPO *Brs3*-expressing neurons and representative expression of channelrhodopsin-2 fused to enhanced yellow fluorescent protein (ChR2-EYFP), with 4’,6-diamidino-2-phenylindole (DAPI) counterstaining. ac, anterior commissure. Scale bar, 100 µm. ChR2-expressing mice and mCherry-expressing controls received identical 473 nm photostimulation delivered at 20 Hz using 2 ms pulses. Optical power was set to 1 mW, measured at the distal end of the patch cord before connection to the implanted optical fiber. (B) Real-time place preference assay and representative occupancy heat maps. Entry into one counterbalanced chamber triggered photostimulation, whereas entry into the opposite chamber terminated photostimulation. (C) Percentage of time spent in the photostimulation-paired chamber (n = 9 mice per group). (D) Experimental design for behavioral testing in the absence of floor heating. After a 3 min adaptation period, mice received photostimulation throughout the subsequent 5 min test session. (E, F) Ethograms from individual control (E) and ChR2-expressing (F) mice. Each row represents one mouse. (G) Latency from photostimulation onset to the first jump among ChR2-expressing mice that exhibited jumping. (H–K) Total duration of wall-digging (H), rearing (I), climbing (J), and jumping (K) during the 5 min photostimulation session (n = 9 mice per group). (L) Open-field assay and representative occupancy heat maps during photostimulation. (M, N) Time spent in the center zone (M) and total distance traveled (N) during the 5 min open-field session (n = 9 mice per group). (O) Elevated plus-maze assay and representative occupancy heat maps during photostimulation. (P, Q) Time spent in the open arms (P) and total distance traveled (Q) during the 5 min elevated plus-maze session (n = 9 mice per group). Each point represents one mouse, and bars represent mean ± SEM. Two-tailed unpaired Student’s t-tests were used to compare ChR2-expressing and control mice in (C, H–K, M, N, P, Q). *P < 0.05; **P < 0.01; ***P < 0.001; ****P < 0.0001; ns, not significant.

We next tested whether activation could reproduce actions observed during floor heating. ChR2-expressing and control mice received 5 min of photostimulation in the mesh-equipped chamber while the heating plate remained switched off (Fig 3D). Ethograms showed wall-digging, climbing, and jumping in ChR2-expressing mice (Fig 3E,F). Jumping began after photostimulation onset, with latency varying among mice (Fig 3G). Compared with controls, ChR2-expressing mice spent more time wall-digging (P < 0.05), climbing (P < 0.01), and jumping (P < 0.0001), whereas rearing duration was unchanged (Fig 3H– K). Wall-digging and jumping were not observed in control mice. Event-count analyses supported these effects, showing more wall-digging and jumping events in ChR2-expressing mice. Climbing event number remained unchanged despite the increase in total climbing duration (Fig S2).

In the open field, photostimulation did not alter center time but increased total distance traveled (P < 0.05; Fig 3L–N). In the elevated plus maze, photostimulation increased both open-arm time and total distance traveled (P < 0.05; Fig 3O–Q). Because increased open-arm time occurred together with greater locomotion, it should not be interpreted independently as reduced anxiety-like behavior. These findings instead indicate that activation increased behavioral output across different nonheated contexts. MnPO *Brs3*-expressing neuron activation therefore produced place avoidance and elicited vigorous actions without floor heating while also increasing locomotor activity more broadly.

### Inhibition of MnPO Brs3-Expressing Neurons Redistributes Heat-Evoked Behavior and Weakens Heat-Source Avoidance

To determine whether MnPO *Brs3*-expressing neurons contribute to behavioral responses under strong heating, we expressed Cre-dependent eNpHR3.0 in this population (Fig 4A). Continuous photoinhibition was delivered throughout a 5 min floor heating session with the plate set to 55°C (Fig 4B–D). Compared with controls, photoinhibition reduced climbing and jumping duration (P < 0.01 and P < 0.0001, respectively), increased rearing duration (P < 0.0001), and did not significantly alter wall-digging (Fig 4E–H). Event-count and cumulative-duration analyses showed the same pattern, with increased rearing, reduced climbing and jumping, and no significant change in wall-digging (Fig S3). The increase in rearing and preservation of wall-digging show that photoinhibition redistributed active behavior rather than uniformly suppressing all responses to floor heating.

**Figure 4.**
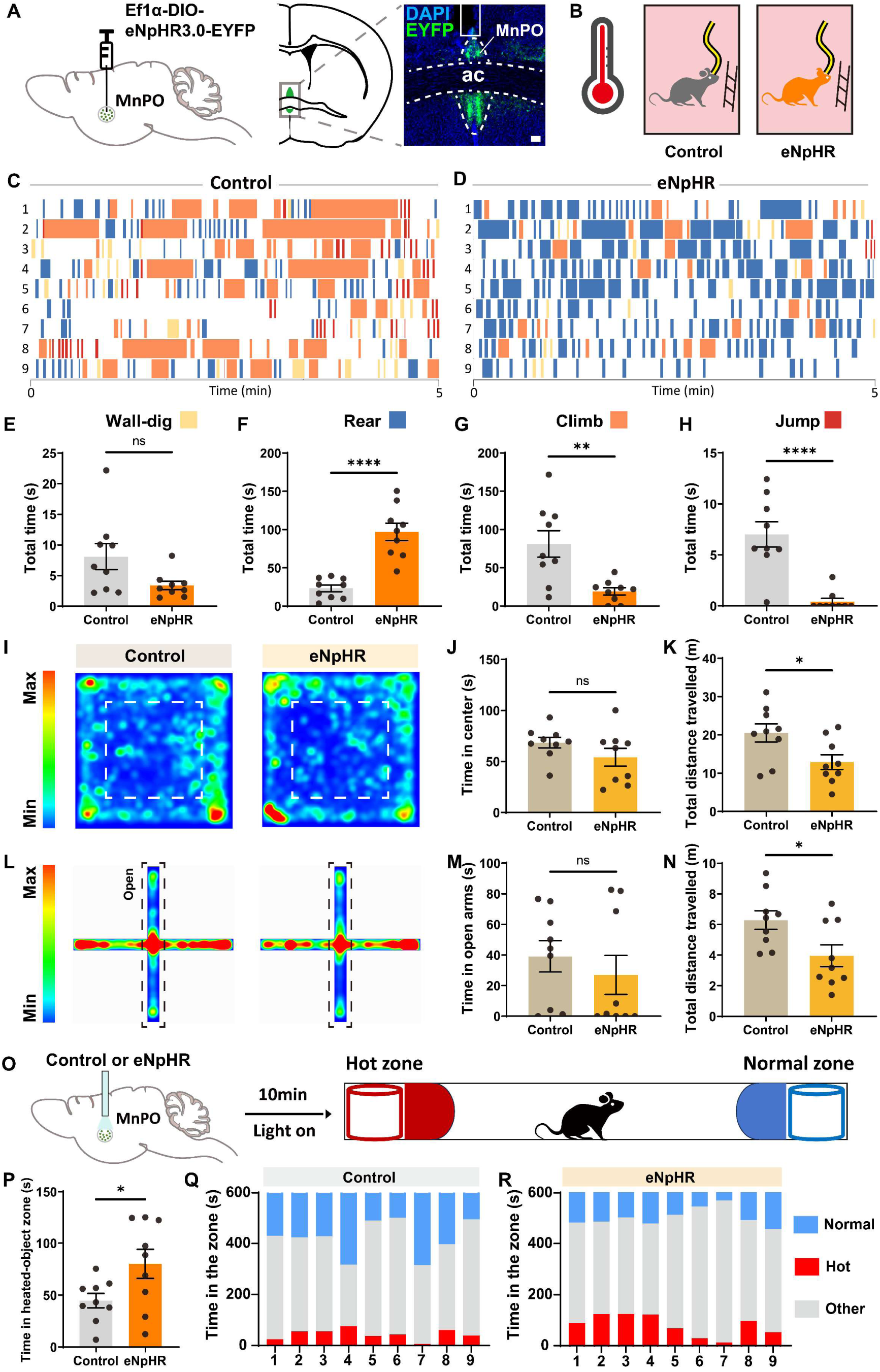
Inhibition of MnPO *Brs3*-expressing neurons shifts heat-evoked behavior away from vigorous actions and weakens heat-source avoidance. (A) Experimental strategy for optogenetic inhibition of MnPO *Brs3*-expressing neurons and representative expression of enhanced Natronomonas pharaonis halorhodopsin 3.0 fused to enhanced yellow fluorescent protein (eNpHR3.0-EYFP), with 4’,6-diamidino-2-phenylindole (DAPI) counterstaining. ac, anterior commissure. Scale bar, 100 µm. eNpHR3.0-expressing mice and mCherry-expressing controls received identical continuous 589 nm illumination. Optical power was set to 3 mW, measured at the distal end of the patch cord before connection to the implanted optical fiber. (B) Experimental design for photoinhibition during floor heating. Mice were tested for 5 min in a chamber with an acrylic floor 5 mm thick positioned above a heating plate set to 55°C. The stated temperature denotes the heating-plate setting. A wall-mounted mesh remained accessible throughout the session, and photoinhibition was delivered continuously during testing. (C, D) Ethograms from individual control (C) and eNpHR3.0-expressing (D) mice. Each row represents one mouse. (E–H) Total duration of wall-digging (E), rearing (F), climbing (G), and jumping (H) during the 5 min floor heating session. (I) Representative open-field occupancy heat maps during photoinhibition. (J, K) Time spent in the center zone (J) and total distance traveled (K) during the 5 min open-field session. (L) Representative elevated plus-maze occupancy heat maps during photoinhibition. (M, N) Time spent in the open arms (M) and total distance traveled (N) during the 5 min elevated plus-maze session. (O) Localized heat-source avoidance assay. Mice explored a corridor containing an object maintained at 55°C at one end and a comparison object maintained at 25°C at the opposite end while receiving continuous photoinhibition for 10 min. The predefined object-adjacent zone extended 100 mm inward from each end of the corridor. (P) Time spent in the zone adjacent to the 55°C object. (Q, R) Occupancy times in the 55°C-object zone, the 25°C-object zone, and the remainder of the corridor for individual control (Q) and eNpHR3.0-expressing (R) mice. For all experiments, n = 9 mice per group. In (E–H, J, K, M, N, P), each point represents one mouse, and bars represent mean ± SEM. In (Q, R), each stacked bar represents one mouse. Two-tailed unpaired Student’s t-tests were used to compare eNpHR3.0-expressing and control mice in (E–H, J, K, M, N, P). *P < 0.05; **P < 0.01; ***P < 0.001; ****P < 0.0001; ns, not significant.

In the open field, photoinhibition reduced total distance traveled without altering center time (P < 0.05; Fig 4I–K). Photoinhibition similarly reduced total distance in the elevated plus maze without altering open-arm time (P < 0.05; Fig 4L–N). The opposing locomotor effects of activation and inhibition indicate that MnPO *Brs3*-expressing neurons influence behavioral activation beyond the immediate response to floor heating. This broader locomotor effect may contribute to the behavioral changes observed during heating but does not fully account for the selective increase in rearing and reduction in climbing and jumping.

We next assessed avoidance of a localized heat source in a corridor containing an object maintained at 55°C at one end and a comparison object maintained at 25°C at the other (Fig 4O). During continuous photoinhibition, eNpHR3.0-expressing mice spent more time than controls in the predefined zone adjacent to the 55°C object (P < 0.05; Fig 4P–R). This increased occupancy is consistent with weaker avoidance of the localized heat source. Because photoinhibition also reduced locomotion, altered exploration of the corridor may have contributed to this effect. Nevertheless, the convergence of weaker heat-source avoidance with activation-induced place avoidance and the selective redistribution of behavior during floor heating supports a role for MnPO *Brs3*-expressing neurons in organizing responses to an aversive thermal challenge. Bidirectional manipulation indicates that this population influences both the vigor and pattern of behavioral responding rather than specifying a single defensive action.

### MnPO Brs3-Expressing Neurons Show Distributed Axonal Labeling

To examine the anatomical distribution of axons arising from MnPO *Brs3*-expressing neurons, we expressed Cre-dependent mCherry in the MnPO and surveyed mCherry-positive fibers across the brain (Fig S4A). Axonal labeling was detected in cortical, striatal, septal, hypothalamic, amygdalar, hippocampal, and midbrain regions (n = 5 mice; Fig S4B). Relatively prominent labeling was found in the ventral subiculum, medial part of the anterior olfactory nucleus, lateral septum, nucleus accumbens core, and zona incerta. Representative sections also showed labeled fibers in the medial orbital cortex, stria terminalis, ventromedial hypothalamus, posterior basolateral amygdala, and posteromedial amygdalohippocampal area (Fig S4C–I). Selected labeled regions are summarized in Fig S4J.

This distributed axonal pattern provides candidate anatomical routes through which MnPO *Brs3*-expressing neurons could influence motivational state, behavioral activation, and thermal defense (Lefler et al., 2020; Silva and McNaughton, 2019; Wang et al., 2015). Axonal labeling alone, however, does not establish synaptic connectivity or assign a specific function to any pathway. Projection-defined recording and manipulation will be needed to determine whether place avoidance, vigorous behavioral output, and heat-source avoidance depend on shared or distinct downstream targets.

Several limitations define the scope of the conclusions. The heating-plate settings were not direct measurements of temperature at the acrylic surface or paw interface. The 55°C setting therefore denotes the strongest heating condition tested rather than a measured paw temperature or defined nociceptive threshold. Core body temperature and autonomic heat-defense responses were not monitored, leaving their relationship to the observed behavior unresolved. Viral expression was centered in the MnPO but extended across dorsal and ventral portions of the nucleus and may have included cells near its anatomical boundaries. The conclusions consequently apply to the MnPO-centered *Brs3*-expressing population accessed by the present targeting strategy. Finally, only adult male mice were studied, and whether the same circuit function is present in females remains to be determined.

Within these boundaries, the results identify MnPO *Brs3*-expressing neurons as a population recruited during strong floor heating that contributes to the transition toward vigorous defensive actions. Their effects on place avoidance, locomotor output, the distribution of heat-evoked behavior, and localized heat-source avoidance place this population at the interface between thermal challenge and behavioral activation. MnPO *Brs3*-expressing neurons therefore shape heat-evoked defensive escalation without specifying a single escape action (Fig 5).

**Figure 5.**
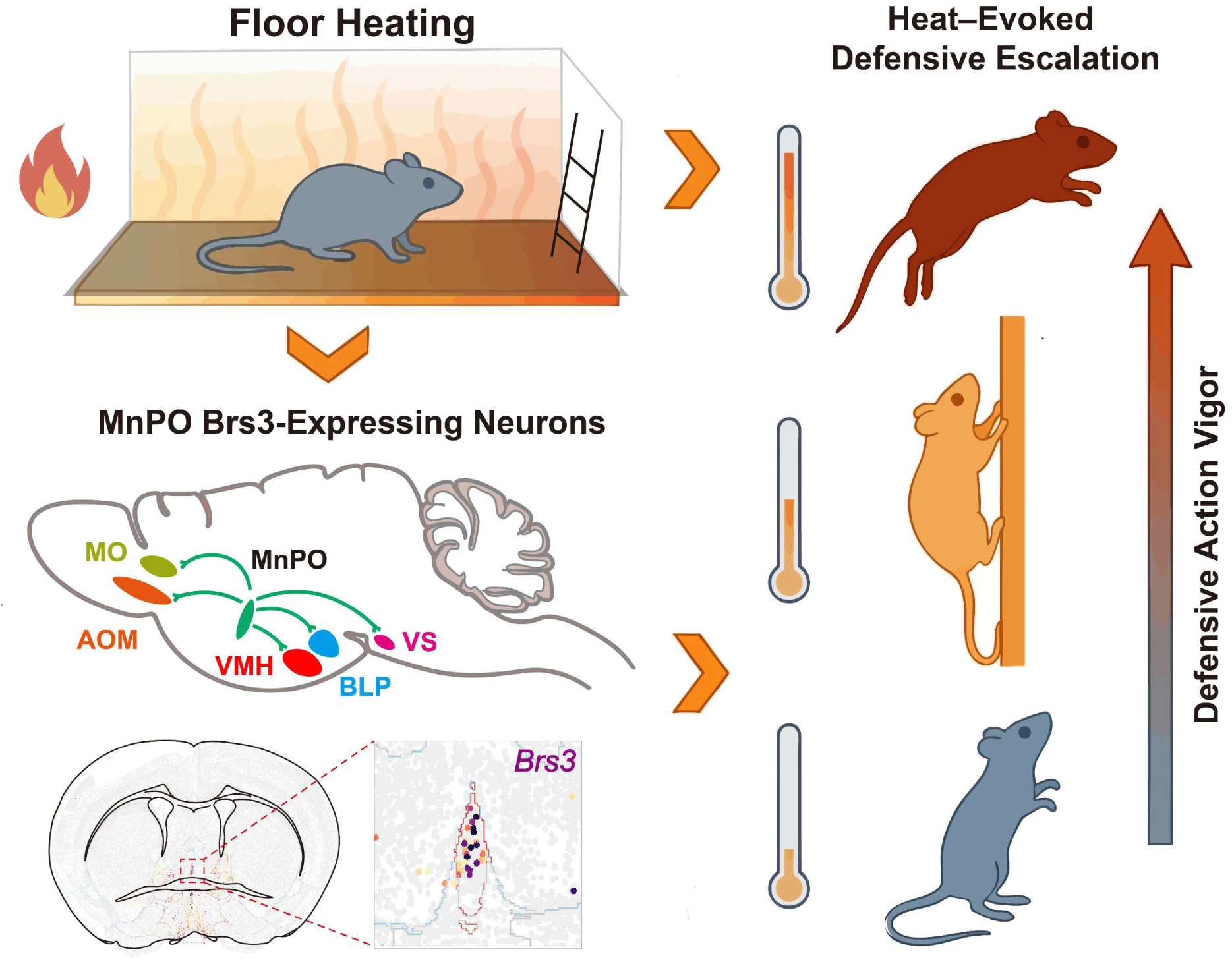
MnPO *Brs3*-expressing neurons shape heat-evoked defensive escalation. As the heating-plate setting increases, mice exposed to floor heating through an acrylic surface, with continuous access to a wall-mounted mesh physically separated from the heating plate, shift their behavioral repertoire from rearing toward more vigorous actions, including wall-digging, climbing, and jumping. MnPO *Brs3*- expressing neurons show increasing recruitment under stronger floor heating and exhibit distributed axonal labeling in regions including the medial orbital cortex (MO), medial part of the anterior olfactory nucleus (AOM), ventromedial hypothalamus (VMH), posterior basolateral amygdala (BLP), and ventral subiculum (VS). The coronal schematic and inset illustrate the spatial distribution of *Brs3*-expressing cells within the MnPO based on publicly available MERFISH data from the Allen Brain Cell Atlas. Activation of MnPO *Brs3*-expressing neurons produces place avoidance and elicits vigorous actions in the absence of floor heating, whereas inhibition shifts heat-evoked behavior toward rearing, reduces climbing and jumping, and weakens avoidance of a localized heat source.

## Methods and Materials

### Animals and ethical approval

Adult male C57BL/6JGpt mice and *Brs3*-P2A-iCre knock-in mice (C57BL/6JGpt-*Brs3*/Gpt; GemPharmatech, strain T006946) were 3 to 4 months old when experiments began. C57BL/6JGpt mice were used for regional c-Fos mapping, whereas *Brs3*-P2A-iCre mice were used for Cre-dependent c-Fos mapping, fiber photometry, optogenetic manipulation, and anterograde axonal labeling. Mice were group-housed at 22 to 24°C under a 12 h light and 12 h dark cycle, with lights on at 08:00, and had unrestricted access to food and water. Animals were allowed to acclimate to the housing facility for at least 1 week before experimental procedures. Only male mice were included, and sex was not evaluated as a biological variable.

The *Brs3*-P2A-iCre genotype was verified by polymerase chain reaction using the forward primer 5’TTCAGTAATTCCTGCGTGAACCC-3’ and the reverse primer 5’TTCCAGGTGTGTTCAGAGAAGGC-3’. The targeted allele generated a 449 bp amplicon (Fig S1). All procedures were approved by the Institutional Animal Care and Use Committee of Southeast University and conducted in accordance with institutional and national guidelines for the care and use of laboratory animals.

### Public MERFISH data

The anatomical distribution of *Brs3*-expressing cells in the MnPO was examined using publicly available multiplexed error-robust fluorescence in situ hybridization data from the Allen Brain Cell Atlas. Coronal sections spanning Bregma +0.59 to −0.06 mm were surveyed, and a section at Bregma +0.13 mm was selected for representative visualization. Cells expressing *Brs3* within the anatomically defined MnPO were displayed to provide spatial context for the viral targeting experiments. The public dataset was used for visualization and was not included in inferential statistical analyses.

### Floor-heating apparatus

Floor heating was delivered in a transparent test chamber whose floor consisted of an acrylic sheet 5 mm thick positioned above a temperature-controlled heating plate. The acrylic sheet separated the mouse from direct contact with the plate. A stainless steel mesh insert was mounted vertically on the chamber wall and was physically separated from the heating plate. The mesh remained accessible throughout testing and provided an alternative surface onto which mice could climb.

The heating plate was set to 25, 35, 45, or 55°C and maintained at the selected setting throughout each session. All temperatures reported for this assay refer to heating-plate settings and not to temperatures measured at the acrylic surface or paw interface. Each session lasted 5 min and was continuously video-recorded. The same acrylic floor and wall-mounted mesh configuration was used for behavioral testing and regional c-Fos mapping. For fiber photometry, the plate-controlled zone incorporated the same acrylic interface.

### Stereotaxic surgery and viral delivery

Mice were anesthetized with 1 to 2% isoflurane in oxygen delivered at 1 L/min and secured in a stereotaxic frame (model 68001; RWD Life Science). Body temperature was maintained at 37°C using a homeothermic blanket. After craniotomy, viral vectors were delivered to the midline MnPO at the following coordinates: anteroposterior +0.14 mm and mediolateral 0.00 mm relative to Bregma, and dorsoventral −4.00 mm from the dura.

Cre-dependent adeno-associated viral vectors carrying a double-floxed inverted open reading frame under the elongation factor 1α promoter were obtained from BrainVTA Technology. The constructs used were AAV-Ef1α-DIO-mCherry, AAV-Ef1α-DIO-GCaMP6f, AAV-Ef1α-DIO-ChR2(H134R)-EYFP, and AAV-Ef1α-DIO-eNpHR3.0-EYFP. AAV2/9-Ef1α-DIO-mCherry was used for anterograde axonal labeling.

A total volume of 90 nL was delivered as a single midline injection over 3 min at 30 nL/min through a glass micropipette connected to a microinjection pump (KD Scientific). The micropipette was left in place for 10 min after infusion and then withdrawn slowly. For fiber photometry and optogenetic experiments, an optical fiber with a core diameter of 200 µm and a numerical aperture of 0.37 was implanted 0.2 mm above the injection center. Mice received carprofen at 5 mg/kg subcutaneously after surgery and were allowed to recover for at least 3 weeks before behavioral testing.

### Anatomical verification and inclusion criteria

Viral expression and optical-fiber placement were examined histologically after completion of the experiments. Expression sites were reconstructed on anatomically matched coronal sections using the MnPO, anterior commissure, and adjacent preoptic structures as landmarks. Animals were included when the center of viral expression was located within the MnPO and, for optical experiments, the fiber tip was positioned above the labeled territory. Animals were excluded when expression was centered outside the MnPO, was predominantly located in an adjacent preoptic region, or when the fiber tip did not overlie the labeled territory. Anatomical inclusion was assessed without reference to behavioral or photometry outcomes.

The injection coordinate was selected to center viral expression in the MnPO near the anterior commissure. Because labeling could encompass dorsal and ventral portions of the MnPO and extend to tissue near its anatomical boundaries, the term MnPO *Brs3*-expressing neurons refers to the MnPO-centered population accessed by this targeting strategy.

### Histology and immunostaining

Mice were deeply anesthetized with isoflurane and transcardially perfused with ice-cold phosphate-buffered saline, followed by 4% paraformaldehyde in phosphate-buffered saline. Brains were postfixed overnight in 4% paraformaldehyde at 4°C and cryoprotected in 30% sucrose. Tissue was embedded in Tissue-Tek O.C.T. compound (Sakura), and 40 µm coronal sections were prepared using a CM1950 cryostat (Leica). Free-floating sections were collected serially and stored in cryoprotectant at −20°C until processing.

For c-Fos immunostaining, sections were washed in phosphate-buffered saline and blocked for 2 h at room temperature in phosphate-buffered saline containing 5% normal goat serum and 0.3% Triton X-100. Sections were incubated overnight at 4°C with rabbit anti-c-Fos antibody diluted 1:1,000 (catalog 2250S; Cell Signaling Technology). After washing, sections were incubated for 2 h at room temperature with either CoraLite 488-conjugated goat anti-rabbit IgG diluted 1:500 (catalog SA00013-2; Proteintech) or Cy3-conjugated goat anti-rabbit IgG diluted 1:500 (catalog SA00009-2; Proteintech), depending on the fluorophore combination required for the experiment. Sections were counterstained with 4’,6-diamidino-2-phenylindole and mounted in antifade medium. Images were acquired using a Leica DM5000B microscope or an Olympus VS200 slide scanner.

### c-Fos activity mapping

For regional c-Fos mapping, C57BL/6JGpt mice were habituated to the test chamber for 3 consecutive days. Separate groups were then exposed for 5 min to a heating plate set to 25, 35, 45, or 55°C. Mice were perfused 90 min after the beginning of the floor heating session.

To assess recruitment of MnPO *Brs3*-expressing neurons, AAV-Ef1α-DIO-mCherry was delivered to the MnPO of *Brs3*-P2A-iCre mice. After 28 days of viral expression, separate groups were exposed for 5 min to a heating plate set to either 25 or 55°C and perfused 90 min after the beginning of the session. Sections were subsequently processed for c-Fos immunostaining.

For regional mapping, c-Fos-positive nuclei were quantified within an anatomically defined MnPO region of interest at matched rostrocaudal levels. For Cre-dependent mapping, recruitment of the labeled population was expressed as the number of mCherry-positive cells containing a c-Fos-positive nucleus divided by the total number of mCherry-positive cells within the region of interest. Regions of interest were defined using the anterior commissure and surrounding preoptic structures as anatomical landmarks. Region boundaries were assigned without reference to c-Fos signal intensity and were applied consistently to anatomically matched sections across conditions.

Images were analyzed in Fiji/ImageJ version 1.54f using consistent background subtraction and intensity thresholds within each experiment. c-Fos-positive nuclei were identified using the Analyze Particles function with an area range of 50 to 200 µm² and a circularity range of 0.5 to 1.0. Nine anatomically matched sections from three mice were analyzed for each condition. Sections from the same mouse were retained as nested observations and were not considered independent biological replicates.

### Fiber Photometry

AAV-Ef1α-DIO-GCaMP6f was delivered to the MnPO of *Brs3*-P2A-iCre mice, and an optical fiber was implanted above the labeled territory. Population calcium signals were recorded using a dual-wavelength fiber-photometry system (R820; RWD Life Science). During recording, mice moved voluntarily between an unheated comparison zone and a test zone whose acrylic floor was positioned above a temperature-controlled plate. The plate beneath the test zone was set to 25, 35, 45, or 55°C.

GCaMP6f was excited at 470 nm, and a 410 nm reference channel was acquired to correct signal components shared with movement and photobleaching. Light power at the fiber tip was maintained at 30–50 µW for each wavelength, and fluorescence was sampled at 40 Hz. The 410 nm signal was fitted to the 470 nm signal by least-squares linear regression, and the fitted reference component was subtracted from the 470 nm signal. Relative fluorescence change was calculated as ΔF/F = (F_corrected − F_baseline)/F_baseline, where F_baseline was the mean corrected fluorescence during the 2 s preceding each transition.

Peri-event traces were extracted from 5 s before to 5 s after entry into the plate-controlled zone or return to the comparison zone. Event-associated changes in ΔF/F were quantified by calculating the area under the curve during the 0 to 5 s interval after each transition, relative to the pre-transition baseline. At each plate setting, each mouse contributed six or seven paired zone-transition trials, with each trial comprising an entry into the plate-controlled zone and a corresponding return to the comparison zone. Events obtained from the same mouse were treated as repeated within-mouse observations. For visualization, fluorescence traces were converted to z scores using the mean and standard deviation of the corresponding recording session.

### Optogenetic Manipulations

For activation experiments, *Brs3*-P2A-iCre mice received AAV-Ef1α-DIO-ChR2(H134R)-EYFP in the MnPO, whereas control mice received AAV-Ef1α-DIO-mCherry. Both groups received 473 nm photostimulation delivered as 2 ms light pulses at 20 Hz. Optical power was set to 1 mW, measured at the distal end of the patch cord before connection to the implanted optical fiber.

For inhibition experiments, *Brs3*-P2A-iCre mice received AAV-Ef1α-DIO-eNpHR3.0-EYFP in the MnPO, whereas control mice received AAV-Ef1α-DIO-mCherry. Both groups received continuous 589 nm illumination at 3 mW, measured at the distal end of the patch cord before connection to the implanted optical fiber.

### Behavioral testing

Behavioral experiments were conducted during the light phase between 09:00 and 17:00. Mice were handled for 30 min daily for 3 consecutive days and acclimated to the testing room for at least 30 min before each experiment. When mice underwent more than one behavioral assay, tests were ordered from lower to higher anticipated stressfulness. Apparatuses were cleaned with 70% ethanol between animals. Mice were randomly assigned to experimental groups, and chamber or stimulation side was counterbalanced where applicable. Sessions were video-recorded and analyzed offline by experimenters blinded to experimental group.

### Floor-heating assay

After 3 days of habituation to the test chamber, separate groups of mice were tested for 5 min with the heating plate set to 25, 35, 45, or 55°C. The selected setting remained constant throughout the session. The wall-mounted mesh insert was available during the entire test. Rearing, wall-digging, climbing, and jumping were scored throughout the session.

### Optogenetic activation without floor heating

ChR2-expressing and control mice were placed in the mesh-equipped chamber and allowed to adapt for 3 min. The heating plate remained switched off. Mice then received 473 nm photostimulation for 5 min, during which rearing, wall-digging, climbing, and jumping were scored.

### Optogenetic inhibition during floor heating

eNpHR3.0-expressing and control mice were tested for 5 min with the heating plate set to 55°C. The wall-mounted mesh remained available throughout the session. Continuous 589 nm illumination was delivered for the entire 5 min test, and rearing, wall-digging, climbing, and jumping were scored offline.

### Behavioral scoring

Manual behavioral scoring was performed using BORIS version 7.9.6. Wall-digging was defined as repetitive forepaw scratching or digging movements directed toward the chamber wall and lasting at least 600 ms. Rearing was defined as standing upright on the hindlimbs, with the forelimbs free or supported against the chamber wall, without sustained upward displacement. Climbing was defined as coordinated upward movement along the wall-mounted mesh involving both forelimbs and hindlimbs. Jumping was defined as a vertical or lateral leap in which all four paws left the floor simultaneously. The duration and number of events for each behavior were quantified over the 5 min session.

### Real-time place preference

The motivational consequences of optogenetic activation were assessed using a two-chamber real-time place preference assay. During a single 5 min session, entry into one counterbalanced chamber triggered 473 nm photostimulation, whereas entry into the opposite chamber terminated photostimulation.

Photostimulation was delivered at 20 Hz using 2 ms pulses, with optical power set to 1 mW at the distal end of the patch cord before connection to the implanted optical fiber. ChR2-expressing and control mice received the same chamber-contingent stimulation protocol. The prespecified outcome was the percentage of the session spent in the photostimulation-paired chamber.

### Open-field test

Mice were placed in the center of a 50 × 50 × 50 cm arena and allowed to explore for 5 min. Illumination at the arena floor was approximately 100 lux. The center zone measured 30 × 30 cm. Time spent in the center zone and total distance traveled were quantified while the corresponding activation or inhibition protocol was delivered throughout the session.

### Elevated plus-maze test

The elevated plus maze consisted of two open arms and two enclosed arms, each measuring 30 × 5.5 cm, elevated 60 cm above the floor. Mice were placed on the central platform facing a closed arm and allowed to explore for 5 min. Illumination on the open arms was approximately 20 lux. Time spent in the open arms and total distance traveled were quantified while the corresponding optogenetic protocol was delivered throughout the session. Open-field and elevated plus-maze behavior was tracked using ANY-maze version 7.4.

### Localized heat-source avoidance

eNpHR3.0-expressing and control mice were tested in a corridor-shaped arena measuring 700 mm long and 50 mm wide. An object maintained at 55°C was positioned at one end, and a comparison object maintained at 25°C was positioned at the opposite end. Before testing, the 100 mm region extending inward from each object was defined as the corresponding object-adjacent zone. Mice explored the corridor for 10 min while receiving continuous 589 nm illumination. The primary outcome was the time spent in the predefined zone adjacent to the 55°C object. Time spent in the 25°C object-adjacent zone and the remainder of the corridor was also quantified.

### Anterograde axonal labeling

AAV2/9-Ef1α-DIO-mCherry was delivered to the MnPO of *Brs3*-P2A-iCre mice. After 3 weeks of expression, brains were processed for histology, and mCherry-labeled axons were surveyed across anatomically defined regions in five mice. For each mouse, labeling assigned to an individual region was divided by the summed signal across all surveyed regions. The resulting values represent the relative distribution of axonal labeling among the regions included in the analysis.

## Statistical Analysis

Statistical analyses were performed using GraphPad Prism version 9.0. Data are presented as mean ± standard error of the mean unless otherwise stated. The experimental unit and definition of n are specified in the corresponding figure legends. Each mouse was treated as one independent biological replicate for between-group behavioral comparisons.

Two independent groups were compared using two-tailed unpaired Student’s t-tests. Predefined event-related changes were compared with zero using two-tailed one-sample t tests. Associations between heating-plate setting and c-Fos labeling or photometry responses were assessed using linear regression. Comparisons between individual heating-plate settings and the 25°C condition were performed using the tests specified in the corresponding figure legends. Tests were two-sided, and P < 0.05 was considered statistically significant.

Histological sections and photometry transitions were sampled repeatedly within individual mice. These observations are explicitly identified as section-level or event-level supporting analyses, and their nesting within mice is reported in the corresponding figures and legends. Sample sizes were based on previous studies and established experimental practice and were not determined by formal power calculations.

## Acknowledgments

We thank the members of the Mu and Xie laboratories for insightful discussions and technical assistance. This work was supported by the National Natural Science Foundation of China (82602934 to M.M.), the General Program of the Natural Science Foundation of Guangdong Province (2026A1515011895 to Z.C.), the Shenzhen Medical Research Fund (A2603009 to Z.C.). The sponsors had no role in study design, data collection, analysis, interpretation, manuscript preparation, or the decision to submit.

## Author Contributions

Conceptualization, M.M.; Methodology, Z.Z., S.H., and M.M.; Investigation, Z.Z., S.H., and M.M.; Visualization, Z.Z. and S.H.; Resources, H.C., Z.C., W.X., and M.M.; Writing— Original Draft, Z.Z. and M.M.; Writing—Review and Editing, all authors; Funding Acquisition, W.X., Z.C., and M.M.; Supervision, M.M.; Project Administration, M.M.

## Data and Materials Availability

All data generated or analyzed during this study are included in the manuscript and its supporting files. The raw data underlying the figures have been provided as Excel files labeled Source Data. Public MERFISH data analyzed in this study are available from the Allen Brain Cell Atlas (https://portal.brain-map.org/atlases-and-data/bkp/abc-atlas). This paper does not report original code. This study did not generate new unique reagents; requests for materials should be directed to Mingdao Mu.

